# The Topology of Evasion: Persistent Homology of Gene Repertoires of Immunity

**DOI:** 10.1101/2025.02.27.640626

**Authors:** Marshall Hampton, Ibrahim Mamudu

## Abstract

Families of genes that are important in adaptive immunity are often large in number and evolve through both mutations and recombination. Extensive recombination produces complex reticulate relationships between genes that require additional analytic tools beyond phylogenetics. We consider several families of genes involved in defense and immune systems from plants, antibiotic resistant bacteria, and trypanosomatid parasites, using the framework of persistent homology from topological data analysis. Some of the gene families considered possess extensive higher-dimensional persistent homological representatives, which may be typical of other gene families participating in adaptive immunity.

## 1. INTRODUCTION

In this work we use a tool of topological data analysis, persistent homology, to examine several remarkable gene families such as the variable surface glycoproteins (VSGs) in the human parasite *Trypanosoma brucei* and leucine-rich repeat (LRR) domain proteins in plants. We also examine simulated sequence evolution to put provide more context for our results.

Most proteins evolve primarily through selection of mutations of DNA sequences within a population. Various processes such as whole genome duplication, or local tandem repetition can produce copies of these sequences, resulting in a unreticulated tree of relationships - a gene phylogeny. In many cases only one copy of such a duplication will functionally persist over long evolutionary periods, but sometimes the duplications will acquire new functions (neofunctionalization) and create a novel protein family. In our species zinc finger proteins are one such family with many hundreds of members. For example, the C2H2 zinc finger subfamily alone has over 700 members in the human proteome [10], which can selectively bind particular DNA and RNA sequences and mediate interactions between other proteins.

Recombination between DNA sequences can result in variant spliced versions of proteins within a family, after which a simple tree phylogeny cannot adequately describe their evolution. Extreme cases of this can arise in protein families in which fast sequence variation provides a strong evolutionary advantage such as in immune defense and evasion. There are a wide variety of proteins involved in immune systems of living creatures, providing many interesting examples. The VSGs and other surface proteins of parasitic trypanosomes are one of the most extreme cases, in which thousands of genes and partial gene sequences rapidly recombine over the course of a single infection to provide variant antigens to evade human or other mammalian immune defenses.

### 1.1. Topological Data Analysis

In this work we will focus on the persistent homology of evolutionary distances of gene families. There is an excellent survey of topological data analysis focused on genomic applications by Rabadán and Blumberg [38]; we also recommend the more general articles [19, 8, 34, 31] and the text [9].

In topology the term homology refers to equivalence classes of sets of geometric objects such as simplices - e.g. points, line segments, triangles, and tetrahedra in dimensions zero to four. A one-dimensional loop is homologically trivial, that is, equivalent to an empty set, if it is the boundary of a two-dimensional disk. Persistence in this context means that as we change a cutoff for what are conisidered “close” distances, some generator of the homology persists for an interval of scale lengths. We will use the simplest choice for constructing the persistent homology, the Vietoris-Rips complex, in which simplices are included in a simplicial complex if all of their sub-simplices are already included; in particular edges are included if their distance is below the given cutoff. Because the computation of the Vietoris-Rips only relies on pairwise distances, it can be computed much faster than more sophisticated alternatives such as the Cěch complex. Even so it can be prohibitively slow to compute higher dimensional homology; in this work we have mainly considered the zero- and one-dimensional homologies (connected components and loops). In our examples it appeared that relevant two-dimensional homology was rare; it would be interesting if other genetic examples with higher-dimensional homological structures could be found.

**Figure 1.**
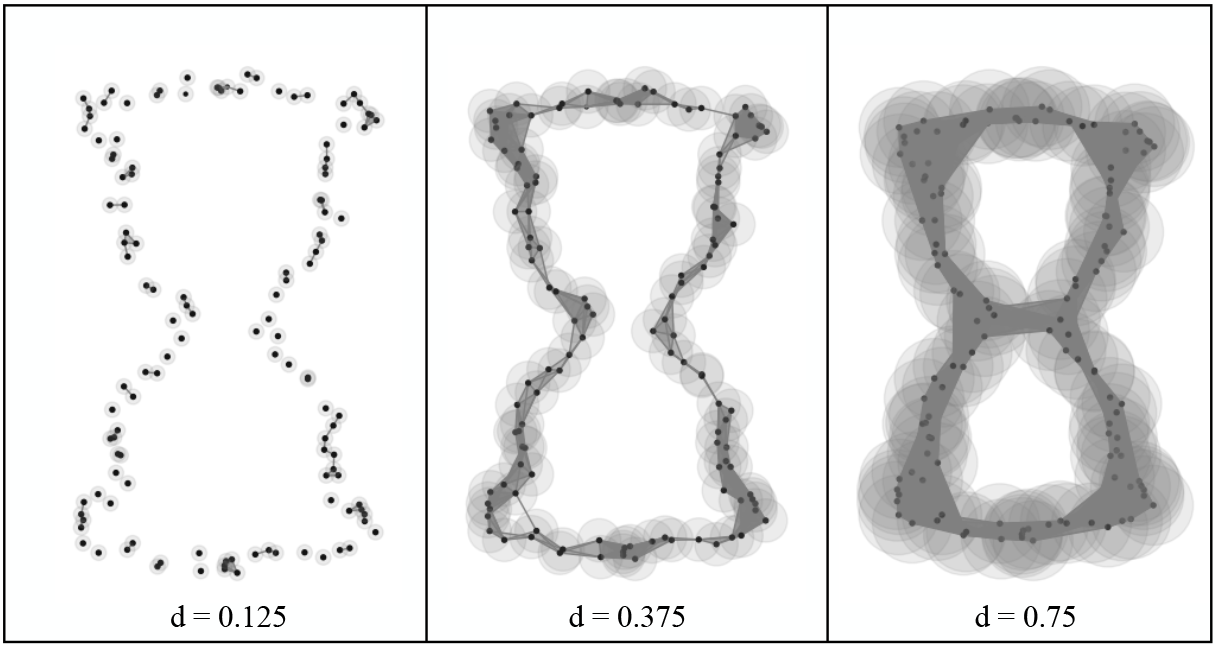
Example of Vietoris-Rips complexes at three different distance cutoffs.

For biological sequence data we use a normalized measure of dissimilarity given by

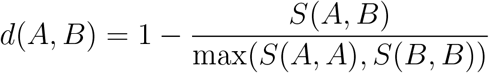

in which *S*(*A, B*) is the alignment score between sequences *A* and *B*. For proteins we compute pairwise alignment scores with NCBI’s Blastp program [2], which uses the BLOSUM62 scoring matrix by default. Sequences which are not successfully aligned by Blastp are given an arbitrary distance larger than 1. This larger distance is not used in any of our analysis as we use a maximum threshold distance of 0.99 in order to avoid artifacts resulting from a lack of alignment. Part of the appeal of persistent homology is that qualitative conclusions should be largely independent of the choice of distance function, especially when comparing sets of data within the same framework.

**Figure 2.**
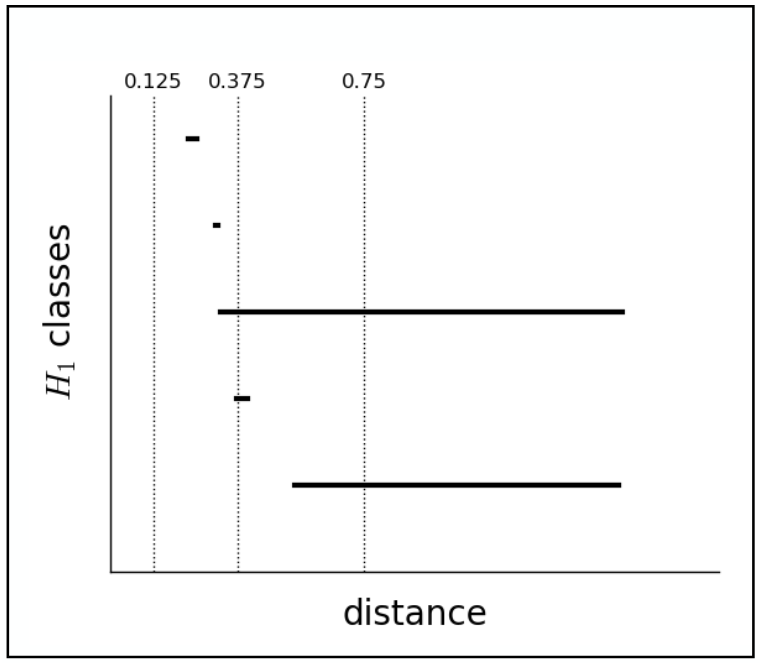
The corresponding *H*_1_ persistent homology barcode, with the three example distances from Figure 1 marked by dotted vertical lines.

The study of the zero-dimensional persistent homology (*H*_0_) of the distances between data points is equivalent to computing a hierarchical clustering of that data [9]. The one-dimensional persistent homology of loops (*H*_1_) has been much less explored in the analysis of biological sequence data. Persistent homology as a tool in data analysis was first described in [18] using somewhat different terminology, and then rediscovered several more times [39, 14]. It has been applied as a tool to study recombination in viruses [11]. Other biological applications of persistent homology and related TDA techniques include classifying breast cancer types [32], plant morphology [25], detecting respiratory pathologies [16], and neuroscience [12, 20, 40].

## 2. Recombination and Persistent Homology

Recombination of DNA is the primary mechanism we would expect to generate significant persistent homology, suggesting that its analysis should provide useful information about the recombinant history of DNA loci.

Using both simulations and extensive genomic data from *Drosophila melanogaster* and *Arabidopsis thaliana*, in [22] it is found that the mean persistence length of the *H*_0_ components and the total number of *H*_1_ generators can together enable a robust estimate of genomic recombination rates. It seems likely that there are more effective statistical approaches, and this is fruitful area for further probabilistic analysis.

In *Escherichia coli*, one estimate is that up to 16% of the genome has been obtained from horizontal gene transfer [33]. In general, bacteria are prone to more horizontal gene transfer than eukaryotes [17]. In this paper we consider two prominent examples: drug-resistant *Staphylococcus* strains, and immune system resistant *Neisseria* species which are responsible for gonorrhea and bacterial meningitis.

From a mathematical perspective the topology of random simplicial complexes has also been studied [24]. However, much remains to be done on the theoretical side for random processes of biological relevance.

In [7], Camara *et al*. used persistent homology to study recombination in the human genome at the nucleotide level.

Based on simulations of recombinant coalescent processes Emmett *et al*. [15] suggested that the expected number of one-dimensional persistent homology features has a distribution of the form

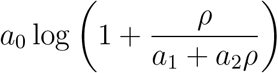

where *a*_0_, *a*_1_, and *a*_2_ are constant and *ρ* is a scaled measure of recombination. For small amounts of recombination this would predict an approximately linear increase in the number of one-dimensional homological features with increasing recombination (*ρ*). We now summarize some of the most relevant biology involved in our study.

### 2.1. Biology of trypanosomes

The Trypanosomatida are a group of eukaryotes within the Euglenozoa which are important because of their unusual biology and disease-causing parasitism. *Trypanosoma brucei* and *Trypanosoma vivax* cause trypanosomiasis and nagana in humans and livestock in Africa, and *T. cruzi* is the cause of Chagas disease in South America.

*T. brucei* are dixenic parasites which alternate between insect hosts (tsetse flies in the genus Glossina) and mammalian hosts including humans. While they are in the mammalian host they predominately live freely in the bloodstream and must constantly evade the adaptive immune response. One of their primary means of defense is the expression of variant surface glycoproteins (VSGs). Only one VSG is primarily expressed at a time [23], but after some time the choice of VSG is switched either by changing the expression site or by ectopic gene conversion. Large numbers of VSG genes and pseudogenes are present near the telomeric ends of the chromosomes as well as smaller numbers in other locations including the minichromosomes. Studies of VSGs historically led to breakthroughs in telomere biology applicable to humans as well [5].

The 3’ end of the mRNA transcripts of expressed VSGs contain a highly conserved sequence [27], the C-terminal domain (CTD), which is associated with the GPI-anchored portion of the protein close to the cell membrane, and is relatively inaccessible to antibody interactions.

We also consider some gene families from *T. cruzi* and *T. vivax. T. vivax* is very similar in lifecycle to *T. brucei* and also possesses a large repertoire of VSG proteins [36], which are quite divergent in sequence from those in *T. brucei*. Unlike *T. brucei*, when in a human host *T. cruzi* parasites eventually enter cells. This means they do not need the elaborate VSG-switching defense of *T. brucei*, but they do need to manipulate the human immune system in other ways. [21]. Their surface proteins also show a remarkable diversity, and provide a good context for comparison with the VSGs of *T. brucei* [30].

### 2.2. Antibiotic and immune system resistant bacteria

*Neisseria* **and** *Staphylococcus*. An important example of reticulated evolution in immune evasion occurs in species of the bacterial genus *Neisseria. N. gonorrhoeae*, the causative agent of the disease gonorrhea, is capable of substantial antigenic variation in its surface proteins including type IV pili proteins, Opa proteins, and sialylated lipooligosaccharides [43, 6, 37, 45]. The type IV pili proteins have a silent (not directly expressed) bank of approximately 20 copies in the *pilS* genomic locus, which recombine with the active *pilE* locus copy. This is very similar in control architecture to the VSG system in trypanosomes.

Similarly, *Neisseria meningitidis*, which is responsible for much of bacterial meningitis and meningococcemia, exhibits unusual genomic variation [42] with several large families of repetitive DNA elements and genes that may play important roles in immune evasion.

We focused our efforts on *N. gonorrhoeae* in this study due to better data availability. The *Staphylococcus* genus contains several species that cause disease in humans, most notably *S. aureus, S. saprophyticus*, and *S. epidermidis*. Drug resistance in *Staphylococcus* has been linked to the exchange of DNA plasmids, particularly in the case of vancomycin resistance. Vancomycin is commonly used to treat methicillin-resistant *S. aureus* (MRSA) infections, but over the last thirty years increasing numbers of vancomycin-resistant strains have arisen. The vancomycin resistance is enabled by vanA operon genes in a transposable element (Tn1546) [3, 28]. It is unclear how exactly the drug resistant version of the operon evolved. It has been found in a variety of other organisms such as the biopesticide *Paenibacillus popilliae* [35]. The resistance operon can be exchanged through plasmids to other pathogenic bacteria such as *Enterococcus faecium* and *Clostridium difficile*, creating huge potential for more antibiotic resistant human diseases. We examined sequences similar to the vanA resistance operon in plasmids of all available species to see if there was continuing recombination.

### 2.3. The immune response in plants and leucine-rich repeat proteins

Plants must defend themselves from a wide variety of pathogens and predators, including fungi, insects, vertebrates, bacteria, and viruses [13]. Among their many methods of defense are classes of proteins capable of recognizing many specific ligands in the extracellular environment, which often include copies of the leucine-rich repeat (LRR) domain. The flexibility of the LRR architecture has resulted in a major diversification of this family in most plant species. Because of this rich variety of function and domains, we chose to examine the LRR family in a selection of 63 high quality plant proteomes.

## 3. Results

### 3.1. Simulations

Adding recombinant events to a branching mutational process greatly increases the number and persistence of one-dimensional generators of homology. In Figure 3 we show *H*_1_ barcode plots of three types of simulations. The first simulation is for a tree of sequences with a stochastic mutational process but no recombination. The second simulation has the same tree geometry (topology and branch lengths), but with recombination between random pairs of sequences at a random homologous subsequence. The final simulation has a much more structured recombination process in which one of four fixed subsequences can be swapped between random pairs. The examples in Figure 3 are representative in that the total length and number of loop classes increase from the first through the third type of simulation.

**Figure 3.**
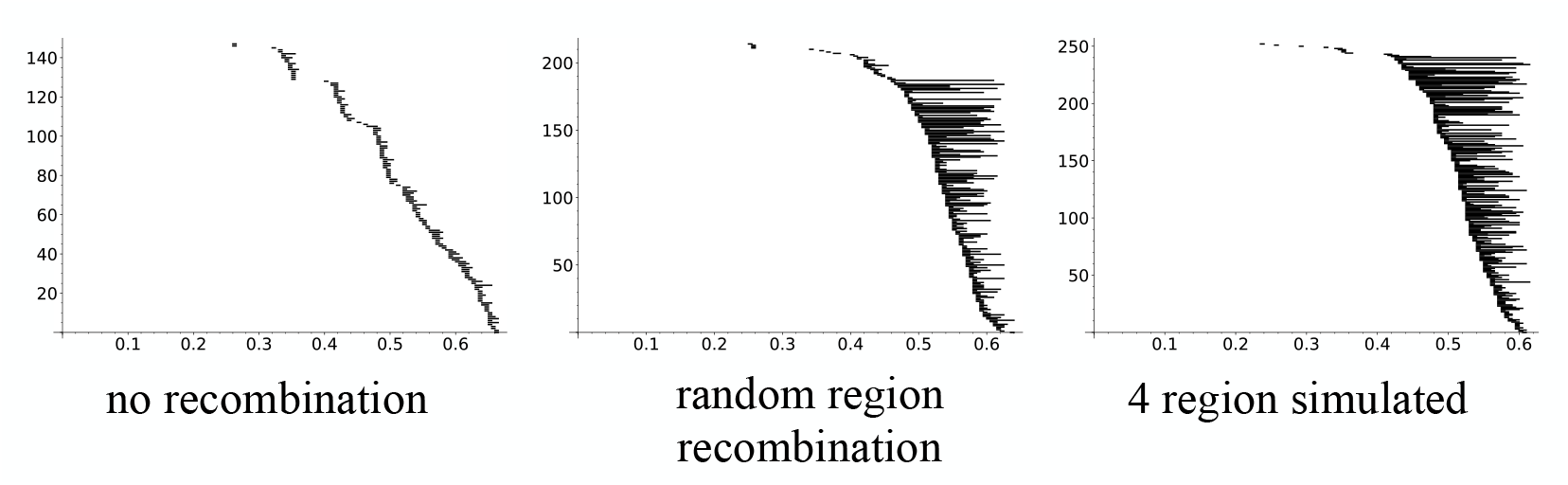
*H*_1_ barcodes for simulated sequence evolution

We believe a great deal of future theoretical work can be done on the statistical signature of different types of recombination processes within the persistent homology data.

### 3.2. Trypanosomal protein families

In *T. cruzi* the three large surface protein families that we examined exhibited very different homological structure, suggesting that the recombination processes within these families may also be quite distinct from one another. The GP82 family had the longest total *H*_1_ length per family member of the three, and may be under the most intense evolutionary pressure to diversify. In contrast the DGF-1 family showed surprisingly little *H*_1_ structure and may be evolving in a more conventional phylogenetic manner.

**Figure 4.**
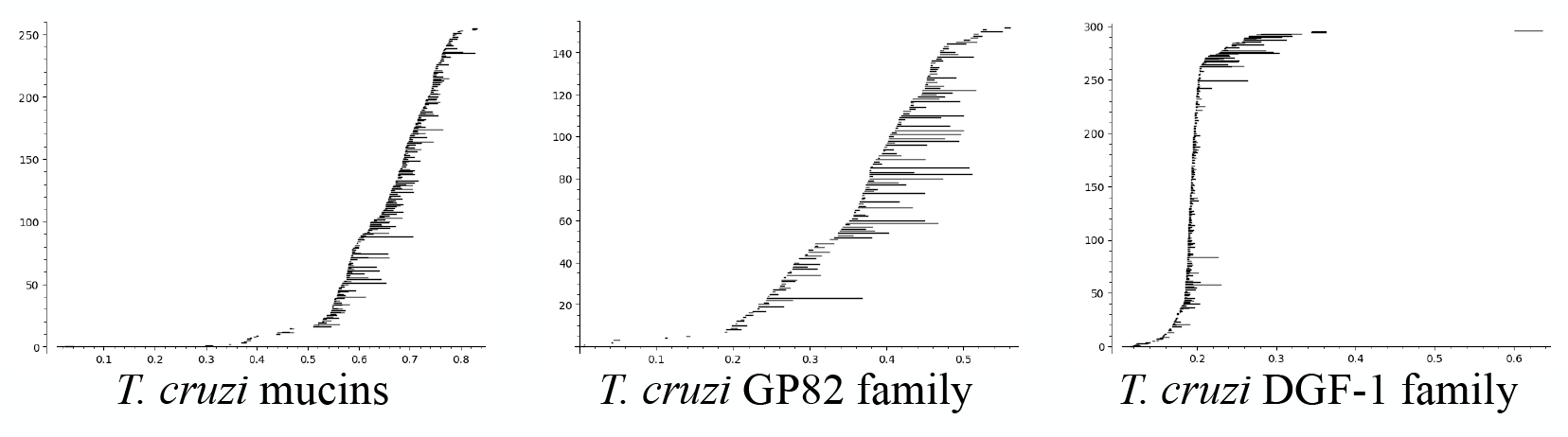
*H*_1_ barcodes for *T. cruzi* protein families

**Figure 5.**
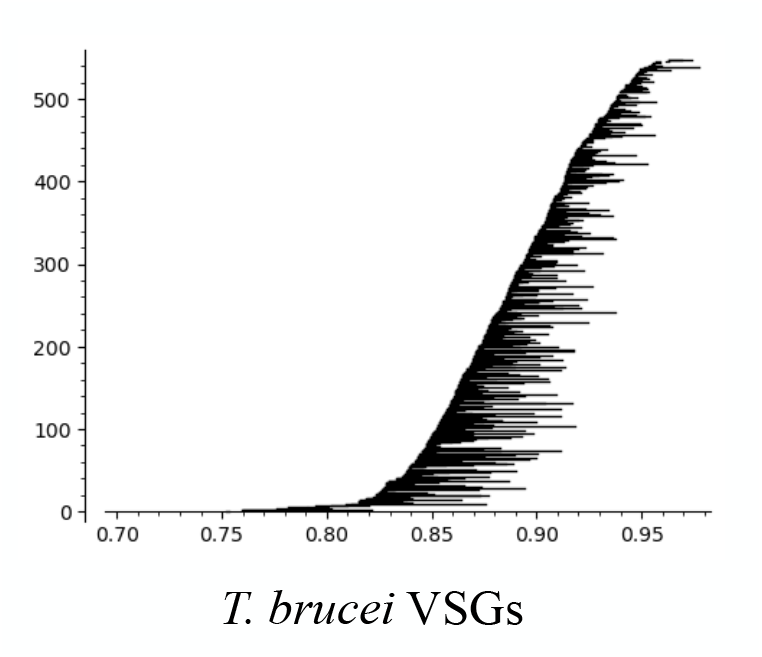
*H*_1_ barcodes for *T. brucei* VSG protein family

As might be expected, the huge VSG family in *T. brucei* possesses a dramatically large amount of persistent loops at a relatively large divergence distance. In order to keep the data comparable to the other protein families considered in this work we limited the number of VSG proteins considered to 512 sequences. A caveat to this analysis is that the large divergence in sequence between the variable portions of the VSG proteins makes reliable alignments difficult. [21]

### 3.3. Bacterial evasion

The pilin proteins of *N. gonorrhoeae* exhibited large amounts of *H*_1_ structure, as shown in Figure 6, providing confirmation that extensive recombination takes place within this family.

**Figure 6.**
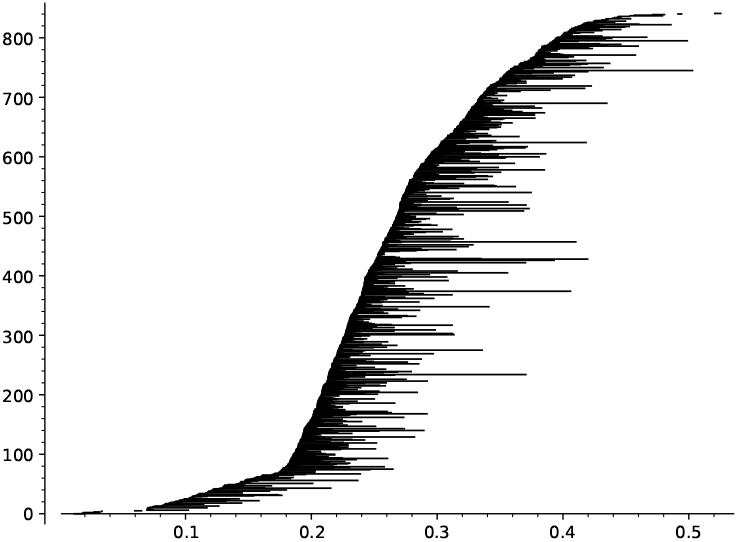
*H*_1_ barcodes for *N. gonorrhoeae* pilin protein family

In contrast we did not find as strong a *H*_1_ signal within the vanA operon sequences taken from the plasmid database PLSDB. However, it seems very possible that this is caused by the relatively small amount of sequences available compared to the vast repertoire present in the global environment.

### 3.4. Plant defense and other proteins

Within the defensive LRR protein family there was a large variation in number and *H*_1_ persistent homology between species. The species with the most *H*_1_ persistent elements, both in number and total persistence length, was the rose gum eucalyptus *Eucalyptus grandis*, followed by the walnut *Juglans regia*. In Figure 7 we show the barcodes for these two species along with those for the model plants *Arabidopsis thaliana* and the black cottonwood *Populus trichocarpa*. The rather extreme difference between these pairs suggest that some species have experienced much more diversification in the LRR family than others, and this may be the result of significantly more recombination within the family in those species.

**Figure 7.**
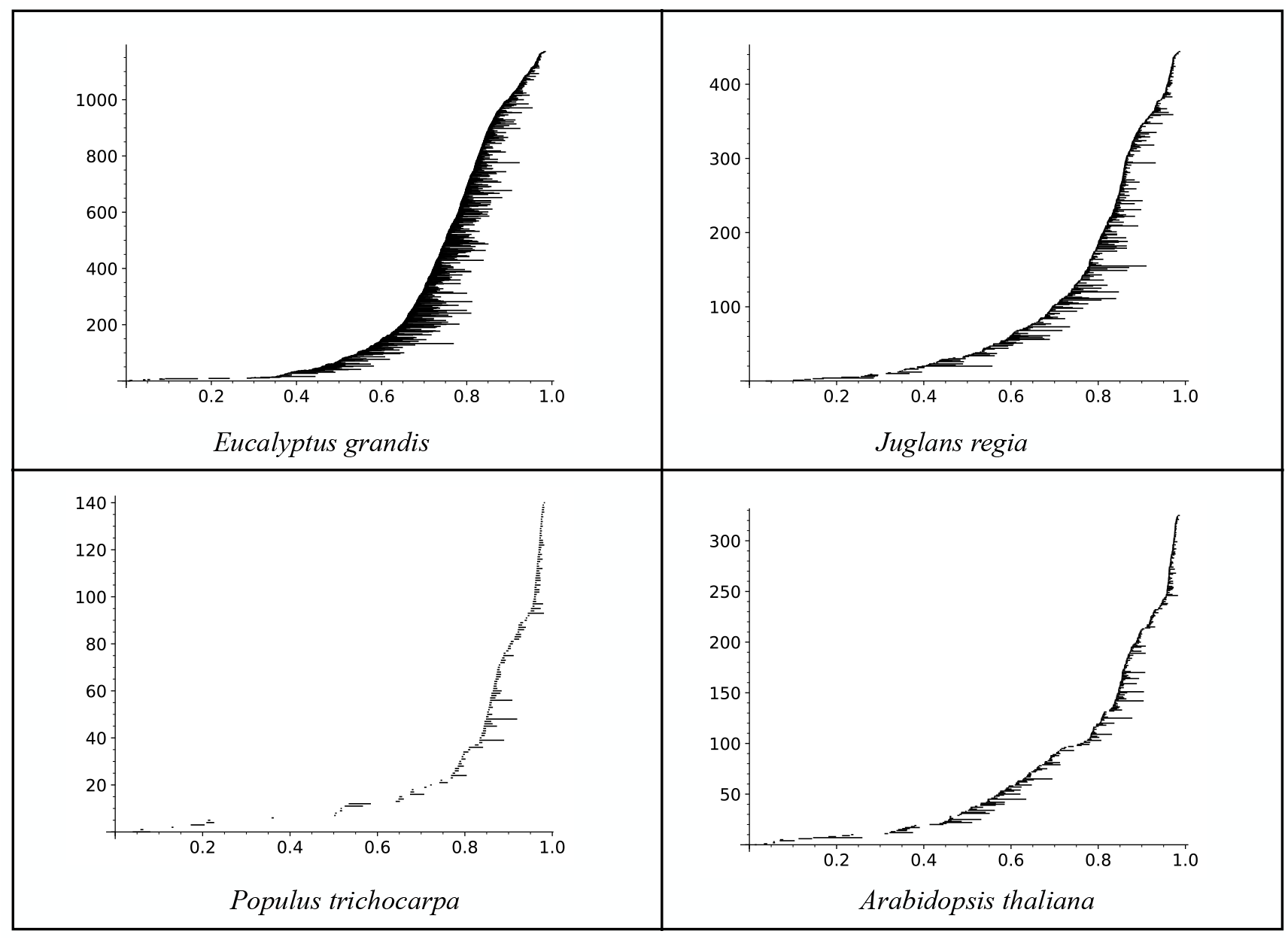
Some plant LRR protein family *H*_1_ barcodes. The horizontal axis is distance, and the vertical is indexing the one-dimensional homology elements. *E. grandis* has the largest total length of *H*_1_ elements.

*Eucalyptus grandis* seems particularly exceptional when we consider the total length of the *H*_1_ persistent homology relative to the number of LRR protein family members in each species. In Figure 8 we show a scatterplot of these statistics. The deviation from the general trend shown by *Eucalyptus grandis* in this context may indicate an unusual recombination process has occurred.

**Figure 8.**
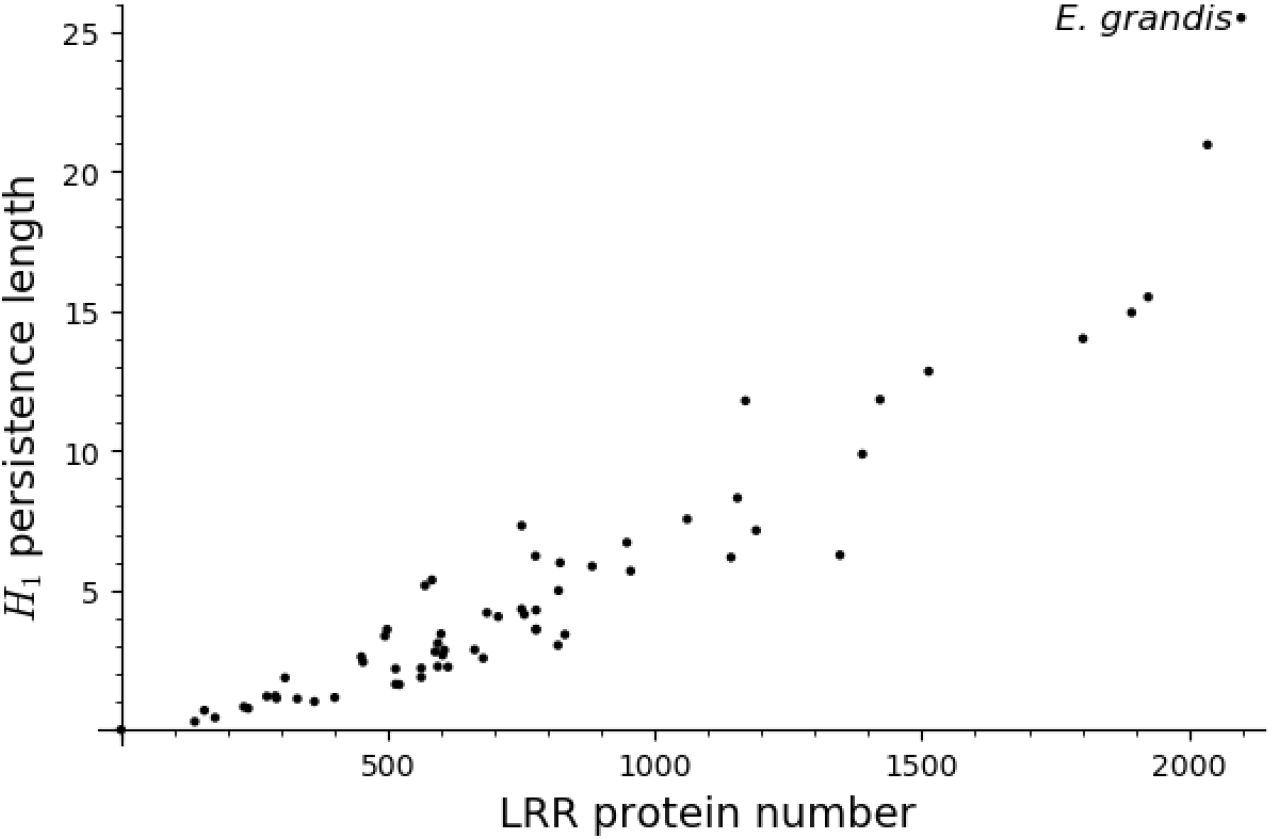
The horizontal axis is the number of LRR protein family members in each of the 63 considered species, and the vertical axis is the total persistent homology length for the one-dimensional homology elements.

**Figure 9.**
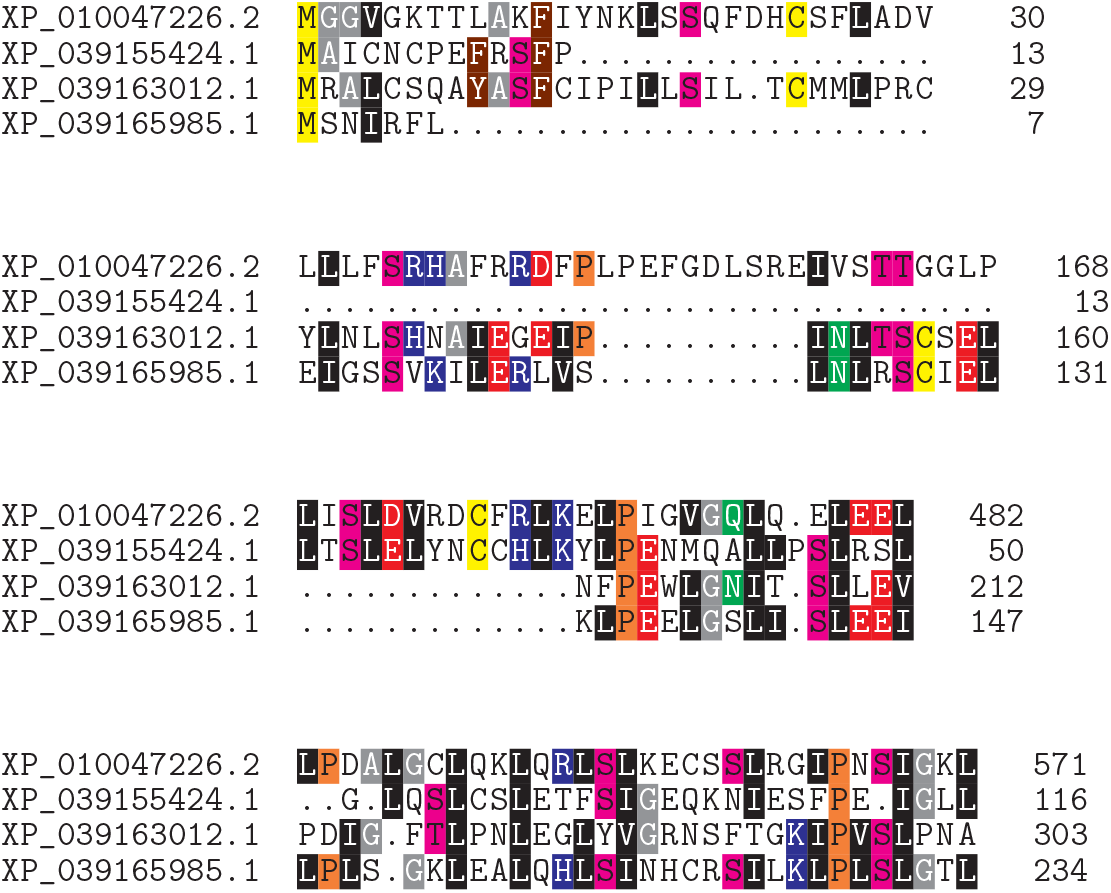
Portions of the alignment of four LRR proteins from *Eucalyptus grandis* that are representatives of the most persistent loop. These sections illustrate the lack of clear phylogenetic relationships among these sequences.

We also computed the zero- and one-dimensional persistent homology of two other large families of proteins within our set of plant proteomes, the P450 cytochrome family and pentatricopeptide repeat (PPR) containing family. There are hundreds to thousands of members of each family in each species considered. While comparable in size to the LRR protein family, the one-dimensional persistent homology was considerably sparser by all measures including total and mean persistence length and number of elements. This was true even for the lycophyte *Selaginella moellendorffii* which has an extremely large number of PPR proteins compared to the other species. This adds some evidence to our hypothesis that recombination is more important for proteins involved in defensive recognition and response processes.

## 4. Methods

All sequences were retrieved from NCBI’s Protein and Nucleotide databases, except Tn1546-matching plasmid sequences which were obtained from the PLSDB [41]. The plant LRR domain proteins were identified using the PFAM domain PF12799 and HMMER version 3.4 [1]. We have used the program Ripser [4] for all of our persistent homology computations. Barcode figures were created in Sage [44] from the Ripser output.

## 5. Conclusion

Overall our results confirm the utility of using persistent homology as a framework to study recombination and evolutionary relationships within protein families. Many other interesting cases of similarly complex reticulate evolution within gene families remain unexplored. To analyze them more theoretical results linking model parameters to persistent homology features are needed. An interesting example that we did not include in this work is the olfactory receptors in mammals and other vertebrates. Most mammals possess a large family of monoallelically expressed olfactory receptors, which must possess some analogous mechanism to the VSG expression control in trypanosomes [26, 29]. However, an important difference between the VSG genes and olfactory genes is that within a particular organism the olfactory receptors expressed must be diverse between cells although within a single cell the expression is monoallelic. The evolutionary pressures in this case are quite different, perhaps in ways that would be detectable using the persistent homology methods outlined here.

